# Increased male mating success in the presence of prey and rivals in a sexually cannibalistic mantis

**DOI:** 10.1101/2021.01.04.425304

**Authors:** Nathan William Burke, Gregory I Holwell

## Abstract

Pre-copulatory sexual cannibalism—or cannibalism without mating—is expected to promote the evolution of male strategies that enhance mating success and reduce the risk of cannibalism, such as preferential mating with feeding females. However, sexual selection on male competitiveness may alter male courtship decisions in the face of cannibalism risk. We investigated the effect of prey availability and rival presence on male mating decisions in the highly cannibalistic Springbok mantis, *Miomantis caffra*. We found that males approached females more rapidly and mated more often in the presence of prey, suggesting that females distracted with foraging may be less of a threat. The presence of a rival also hastened the onset of copulation and led to higher mating success, with very large effects occurring in the presence of both prey and rivals, indicating that intrasexual competition may intensify attraction to foraging females. Taken together, our results suggest that pre-copulatory cannibalism has selected for male preference for foraging females, and that males adjust their mating strategy to both the risk of competition and the threat of cannibalism.

**LAY SUMMARY:** Deciding when to approach a mate is critical for male mantises at risk of being cannibalised. A male might do well to pounce when a female is distracted with prey, but what if a nearby male has the same intention? In the Springbok mantis, we show that males mate faster and with greater success when both prey and a rival are present, suggesting that mating decisions depend on the dual threats of cannibalism and competition.

## INTRODUCTION

Understanding how animals maximise fitness while managing risk is a key focus of behavioural ecology (Magnhagen 1991; Dall and Johnstone 2002; Dall 2010; Mathot et al. 2012). In sexually cannibalistic species—where females consume males prior to, during, or immediately following copulation—males must balance the imperative to mate with the danger of being eaten by females (Elgar 1992). This balancing act is particularly critical in species where cannibalism occurs without mating (i.e., precopulatory cannibalism) because males forfeit all current and future reproductive success if they are cannibalised (Elgar and Schneider 2004).

Males confronted with extreme costs of precopulatory cannibalism are expected to evolve behavioural tactics to avert or avoid female aggression (Parker 1979; Elgar 1992; Schneider 2014). A common tactic emphasising distraction occurs when males preferentially approach feeding females. Female feeding behaviour may be an important trigger for mating initiation because females distracted by prey may be less likely to attack approaching males (Prenter et al. 1994; Fromhage and Schneider 2005). In some orb-weaving spiders, males typically wait for females to feed on prey before entering the web to attempt mating (Fromhage and Schneider 2005). Similarly, some male mantids prefer feeding females (Scardamaglia et al. 2015) and mate with such females more readily (Gemeno and Claramunt 2006). This suggests that females in prey-abundant environments may be less dangerous for males to approach, and therefore the risks of cannibalism in such environments may be reduced. However, the influence of prey abundance on male mating decisions with respect to cannibalism risk is poorly understood.

The presence of competitors could also influence how males interact with cannibalistic females. Since most sexual cannibals are scramble competitors—where males compete to be the first to find females and mate (Emlen and Oring 1977; Thornhill and Alcock 1983; Herberstein et al. 2017)—the presence of male rivals is likely to feature prominently in a male’s calculus of risk (Prokop and Václav 2005). On the one hand, a male may use a competitor as a decoy or ‘sacrificial lamb’, timing his approach to coincide with a female stalking, attacking or eating the competitor. On the other hand, the presence of rivals may intensify a male’s motivation to mate, which may lead to faster mating approaches and increased mating success, as occurs in non-cannibalistic taxa (Simmons 1986; Beani and Turillazzi 1990; Price and Rodd 2006). Such a lack of caution may however lead to a higher incidence of cannibalism due to hasty missteps (Stoltz et al. 2008). If males pay attention to female foraging behaviour, the presence of prey could modulate competition among males in non-additive ways. For example, if males approach feeding females faster than they approach non-feeding females, the presence of a rival may intensify the speed of their approach, resulting in either higher mating success or higher mating failure due to cannibalism. While effects of female feeding behaviour and rival presence have been investigated in isolation, how such factors interact to shape anti-cannibalism behaviours in males is currently unclear.

The Springbok mantis, *Miomantis caffra*, is an excellent system for investigating effects of prey and competitors on male mating decisions in response to the risk of cannibalism. As in many other mantises, males are scramble competitors (Maxwell 1999). However, unlike most cannibalistic species, pre-copulatory cannibalism occurs at an extremely high rate: more than 60% of inter-sexual encounters end in cannibalism without mating, one of the highest known natural rates of cannibalism (Walker and Holwell 2016). Males have limited capacity to assess the likelihood of being attacked since female aggression is not consistent within individuals (Fisher et al. 2020), and is uninfluenced by female body size, condition, or feeding regime (Walker and Holwell 2016). Females are also facultative parthenogens which means they can produce viable offspring asexually without mating, via parthenogenesis (Walker and Holwell 2016). This unique confluence of traits is expected to impose strong selection on males to evolve tactics that, on the one hand, increase their competitive edge against rivals, and, on the other, mitigate the risk of mating failure due to cannibalism.

We conducted a laboratory experiment on *M. caffra* in which the presence of prey and the presence of a rival were simultaneously manipulated. We hypothesised that males would pay attention to female feeding behaviour by approaching females faster and mating more frequently when prey were present. We also suspected that competition would increase the motivation of males to mate, with the fastest onset of sexual contact occurring in the presence of both prey and a rival. Due to the distracting nature of prey, we further predicted that competition would enhance male mating success when prey were present, but would exacerbate the incidence of cannibalism when prey were absent.

## METHODS

To investigate the influence of rival and prey presence on mating and cannibalism, we performed a fully factorial mating experiment consisting of two interacting treatments that manipulated the presence of heterospecific prey and conspecific males. We placed individual adult virgin females that had not yet oviposited (*n* = 76; 19 females per treatment combination) into separate 30 x 30 x 30 cm mesh enclosures containing a bunch of artificial plastic leaves and introduced either one or two adult males (‘rival treatment’), and either 40 house flies or no flies (‘prey treatment’). To prevent confounding due to differences in female satiation levels between the prey treatment groups, females in the ‘prey absent’ group were each housed with 40 flies for the 12 hours immediately prior to the experimental trial, while females in the ‘prey present’ group were provided with 3 flies during this same period. Observations of mating and/or cannibalism were made every hour for 8 hours. The experiment was conducted in several blocks over multiple days due to space constraints. All mantises were obtained as juveniles from numerous locations in Auckland, New Zealand.

To analyse the likelihood of mating and cannibalism, we used generalised linear models (GLMs) with binomial error structures and logit link functions. Mating outcome and cannibalism outcome were treated as separate binary response variables, and rival treatment, prey treatment and their interaction were included as fixed effects. A Cox proportional hazards regression model was used to assess treatment differences in the onset of mating. Latency to mate (measured in hours) was the response variable, with rival treatment, prey treatment, and their interaction fitted as fixed effects. Individuals that did not mate were treated as censored observations.

We used likelihood ratio tests to assess the significance of fixed effects in all models. This was done by removing each fixed effect from a reduced model that had nonsignificant higher-level interactions removed. The significance of interactions was similarly assessed by removing interaction effects from the full model. Trial date was initially included as a categorical covariate but was later excluded from all analyses as no block effects were detected (−8.911 ≤ χ^2^ ≤ 7.656, 0.780 ≤ *P* ≤ 0.865). For effect sizes of mating success, we report standardised mean differences (*d*) and 95% confidence intervals (CI) using the probit transformation for binary response data (Glass et al. 1981). We report hazard ratios (HR) and their 95% CIs for mating onset effect sizes.

## RESULTS

Males initiated mating nearly 4 times quicker when prey were present (Cox model: HR = 3.799, CI = 1.588 to 9.088; analysis of deviance: χ^2^ = 10.359, *P* = 0.001; Figure 1), and 3 times quicker when a rival male was present (Cox model: HR = 2.968, CI = 1.285 to 6.857; analysis of deviance: χ^2^ = 7.123, *P* = 0.008; Figure 1). Speed was enhanced by the presence of both factors. When a rival was present, the addition of prey increased the speed of approach by 4 times (HR = 4.034, CI = 1.446 to 11.260; Figure 1), and when prey were present, the addition of a rival increased the speed of approach by nearly 6 times (HR = 5.587, CI = 1.825 to 17.100; Figure 1). By comparison, when only a rival was present, mating onset was less than twice as fast as when no rival or prey were present (HR = 1.770, CI = 0.423 to 7.410; Figure 1), and the presence of only prey had a near equivalent effect on mating onset as the presence of no rival or prey (HR = 1.322, CI = 0.296 to 5.907; Figure 1). Despite the pattern of interaction suggested by these effect sizes and CIs, there was no significant interaction effect between prey and rival presence in the model (Cox model: HR = 3.068, CI = 0.500 to 18.821; analysis of deviance: χ^2^ = 1.427, *P* = 0.232).

**Figure 1.**
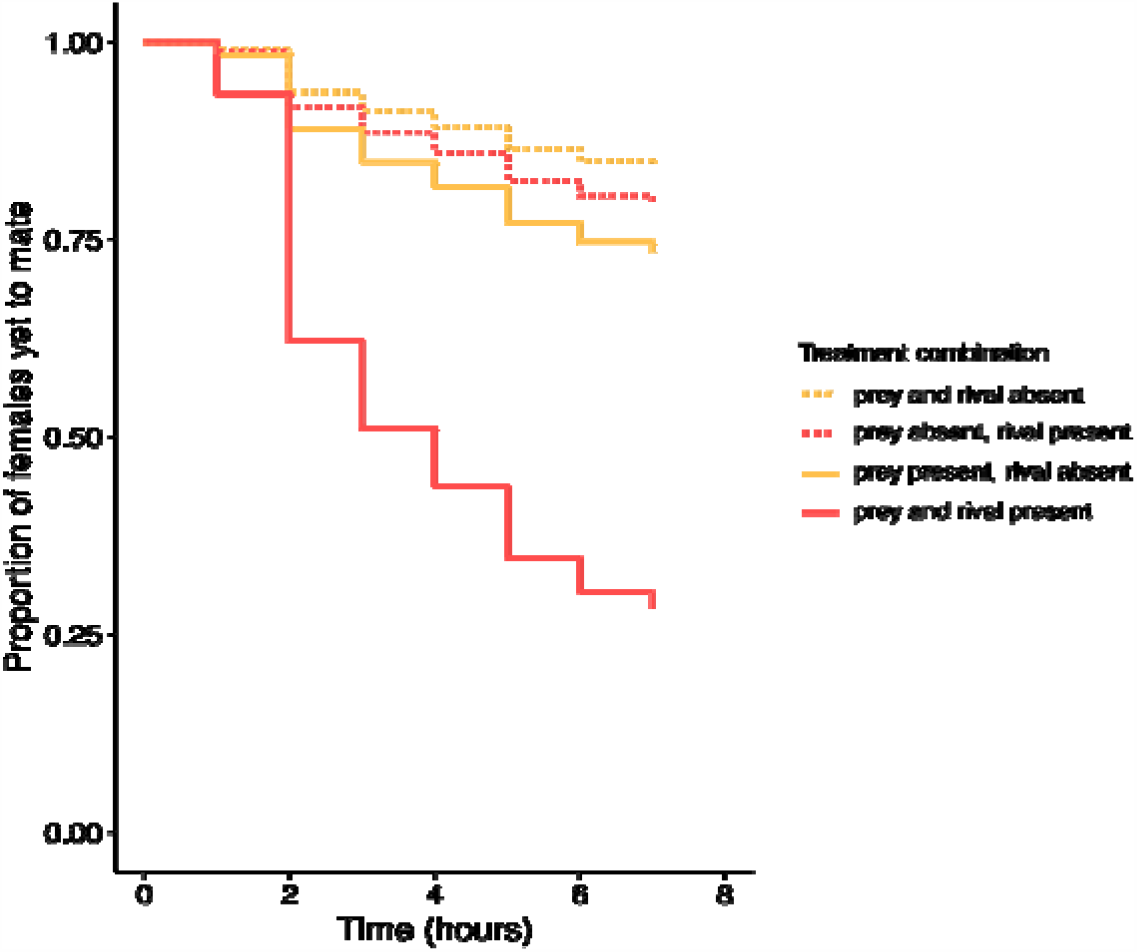
Survival curves showing the proportion of females remaining unmated over the duration of mating trials. Censored observations are not depicted.

The incidence of mating increased 171% in the presence of prey (mating incidence: prey present: 19/38, prey absent: 7/38; *d* = 0.899, CI = 0.289 to 1.510; GLM coefficient = 1.639; analysis of deviance: χ^2^ = −9.460, *P* = 0.002; Figure 2), and 125% in the presence of a male rival (mating incidence: rival present: 18/38, rival absent: 8/38; *d* = 0.739, CI = 0.138 to 1.339; GLM coefficient = 1.391; analysis of deviance: χ^2^ = 6.7585, *P* = 0.009; Figure 2). Prey and rival presence had large interactive effects. The addition of a rival increased mating success by 180% when prey were present (mating incidence: rival with prey: 14/19, no rival with prey: 5/19; *d* = 1.267, CI = 0.409 to 2.125; Figure 2), but only 33% when prey were absent (mating incidence: rival without prey: 4/19, no rival or prey: 3/19; *d* = 0.199, CI = −0.732 to 1.129; Figure 2). Similarly, the presence of prey enhanced mating success by 250% when a rival was present (*d* = 1.257, CI = 0.345 to 2.169; Figure 2), but only 67% when a rival was absent (*d* = 0.370, CI = −0.542 to 1.281; Figure 2). Although the effect sizes and CIs of these pairwise comparisons suggest that prey and rival presence enhanced mating success only when they occurred together, there was no significant interaction effect according to the model (GLM coefficient = 1.707; analysis of deviance: χ^2^ = −2.295, *P* = 0.130).

**Figure 2.**
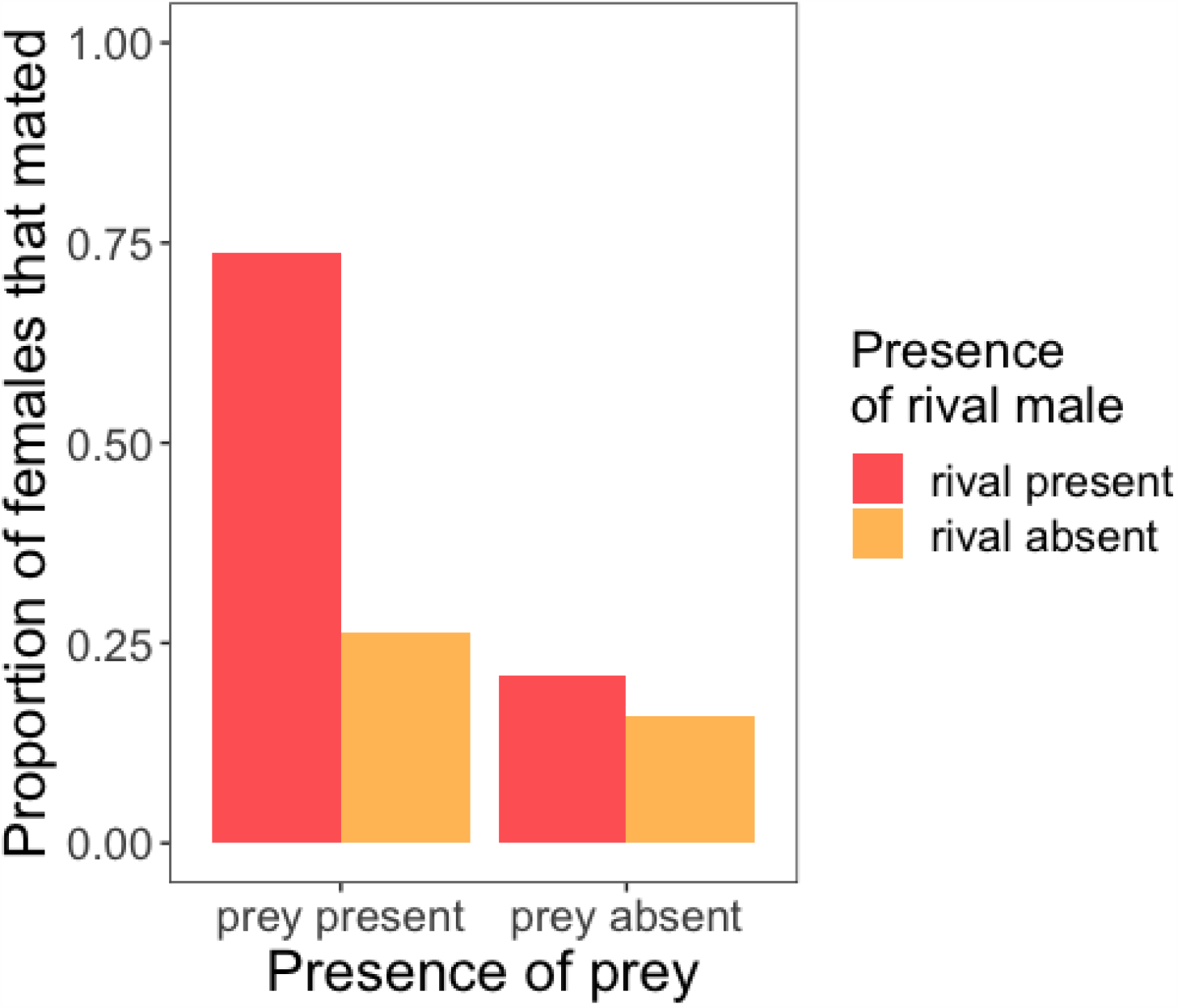
Bar graph showing the proportion of mating trials that ended in mating.

Cannibalism was observed in 2 out of 76 trials, and its occurrence was unaffected by any treatment (−17.676 ≤ GLM coefficient ≤ 35.351; analysis of deviance: −2.827 ≤ χ^2^ ≤ 0, 0.093 ≤ *P* ≤ 1). We observed no incidences of females eating one male while simultaneously copulating with the other.

## DISCUSSION

Our results support the idea that sexually antagonistic selection favours the evolution of male strategies that prevent sexual cannibalism and enhance mating success (Elgar 1992; Schneider and Lubin 1998; Elgar and Schneider 2004). We found that matings were initiated more often and more rapidly in the presence of prey, suggesting that males use female foraging behaviour as a cue to determine the safest time to approach—a common male strategy in sexually cannibalistic taxa (Prenter et al. 1994; Elgar and Fahey 1996; Fromhage and Schneider 2005; Gemeno and Claramunt 2006). Although males were probably attracted to foraging females because of the lower risk they posed, we found no difference in the incidence of cannibalism among treatments. Indeed, rates of cannibalism were significantly lower than those previously reported for this species—a finding likely to be driven by the extremely high satiation levels experienced by females in our experimental design. Nonetheless, as predicted under scramble competition, males responded to the presence of a rival by approaching females faster and more frequently. This effect appeared to depend on the presence of prey: males with both rivals and prey present initiated mating 4 times faster and were 3.5 times more successful than those with only rivals. Taken together, our results illustrate how the risk of mating failure due to precopulatory cannibalism can alter selection on male mating decisions in context-dependent ways, and highlights the potential for sexual selection to modulate antagonistic interactions in sexually cannibalistic taxa.

Our results suggest that males pay attention to the feeding behaviour of females when deciding when to initiate mating. Such a strategy has been suggested for other cannibalistic taxa where males preferentially approach females that are actively handling or eating prey (Prenter et al. 1994; Fromhage and Schneider 2005; Gemeno and Claramunt 2006). However, it is known that males can discriminate between well-fed and hungry females (Barry et al. 2010; Brown et al. 2012). Thus, feeding females could be more attractive not because prey-handling makes them less able to attack but because feeding provides a cue to males that females are becoming satiated and therefore less dangerous (Avigliano et al. 2016). Our experimental design accounted for this possibility by providing females with equivalent access to large numbers of prey immediately before or during trials. This meant that any difference in male approach in the presence versus absence of prey was unlikely to be due to differences in female hunger. We found that matings were initiated more frequently and more quickly when females were actively foraging (i.e., not satiated the previous day), suggesting that, in this case, males assessed the risk of cannibalism using cues associated with female foraging behaviour rather than perceived hunger levels. These results lend strong support to the idea that distraction is an important signal for mating initiation in cannibalistic taxa (Maxwell 1998; Bilde et al. 2006; Uhl et al. 2015; Toft and Albo 2016). However, it is likely that males pay attention to the entire predation sequence when timing their approach, with certain behavioural cues provoking greater response than others (Scardamaglia et al. 2015). Finer-scale observations would be a valuable next step in assessing the relative importance of specific behaviours, such as stalking, striking and feeding, in mating initiation.

Our results are broadly consistent with the prediction that competition to fertilise eggs hastens the onset of mating in protandrous mating systems (Darwin 1871; Thornhill and Alcock 1983; Herberstein et al. 2017). Males in our experiment increased their mating effort in the presence of a rival by initiating matings more rapidly and more frequently. However, rival presence did not alter the incidence of cannibalism. Thus, our results provide no evidence that enhanced motivation to mate under competition causes males to misjudge the risk of cannibalism by approaching females too hastily. This is in contrast to the red back spider in which the presence of a rival increases the incidence of precopulatory cannibalism by causing males to spend less time in premating courtship than is necessary to convince females to mate (Stoltz et al. 2008). However, the low rate of cannibalism in our study was probably due to all females being well satiated rather than because of the presence or absence of male rivals. How competition for females of differing hunger levels affects cannibalism risk therefore remains an open question.

We predicted that the presence of prey could modulate male-male competition for mates if mating tactics are influenced by female feeding behaviour. We found very large effects when both a rival and prey were present, and small effects when only a rival or prey were present, indicating that male mating success is enhanced by the concurrent presence of both cues. This suggests that males may experience a higher risk of competition when females are distracted with foraging, and so approach faster to avoid rivals reaching such females first, resulting in higher mating success. These results highlight the general importance of considering a range of realistic ecological contexts for male anti-cannibalism strategies. Staged experiments that fail to account for important sources of ecological variation (such as the density of prey and competitors) may over-estimate cannibalism rates and under-estimate the effectiveness of male counter adaptations. Further exploration of the influence of prey abundance and quality as well as male abundance in natural populations (e.g., Rabaneda-Bueno et al. 2008) will be necessary for understanding the adaptive significance of sexual cannibalism and its role in sexually antagonistic coevolution.

